# Buruli ulcer surveillance in south-eastern Australian possums: infection status, lesion mapping and internal distribution of *Mycobacterium ulcerans*

**DOI:** 10.1101/2024.05.07.592878

**Authors:** Emma C. Hobbs, Jessica L. Porter, Jean Y.H. Lee, Panayiotis Loukopoulos, Pam Whiteley, Lee F. Skerratt, Timothy P. Stinear, Katherine B. Gibney, Anna L. Meredith

**Affiliations:** Melbourne Veterinary School, Department of Veterinary Biosciences, Faculty of Science, The University of Melbourne, Werribee, Victoria, Australia; Department of Infectious Diseases, Peter Doherty Institute for Infection and Immunity, The University of Melbourne, Parkville, Victoria, Australia; Department of Microbiology and Immunology, Peter Doherty Institute for Infection and Immunity, The University of Melbourne, Parkville, Victoria, Australia; Office of the Dean, Faculty of Natural Sciences, The University of Keele, England, United Kingdom

**Author notes:** TS, KG and AM are Joint Senior Authors.

## Abstract

Buruli ulcer (BU) is a neglected tropical disease of skin and subcutaneous tissues caused by *Mycobacterium ulcerans*. BU-endemic areas are highly focal, and *M. ulcerans* transmission dynamics vary in different settings. In the south-eastern Australian state of Victoria, BU is an endemic vector-borne zoonosis, with mosquitoes and native possums implicated in transmission, and humans as incidental hosts. Despite the importance of possums as wildlife reservoirs of *M. ulcerans*, knowledge of BU in these animals is limited.

Opportunistic necropsy-based and active trap-and-release surveillance studies were conducted in and around Melbourne and Geelong, Victoria, to investigate BU in possums. Demographic data and biological samples were collected, and when present, cutaneous lesions suggestive of BU were mapped. Samples were tested for the presence of *M. ulcerans* DNA by IS2404 qPCR. The final dataset included 26 possums: 20 necropsied; 6 trapped and released. Most possums (77%) were common ringtails from inner Melbourne. Nine possums (eight ringtails, one brushtail) had skin lesions, ranging from single and mild, to multiple and severe, exposing bones and tendons in three cases. *M. ulcerans* was confirmed in 78% (7/9) of clinically affected possums and 65% of possums without lesions (11/17). Possums with moderate and severe disease had widespread systemic internal bacterial dissemination and were shedding *M. ulcerans* in their faeces. The anatomical distribution of cutaneous lesions and PCR positivity of biological samples suggests possums may be contracting BU from bites of *M. ulcerans*-harbouring mosquitoes, traumatic skin wounds, ingestion of an unknown environmental source, and/or during early development in the pouch.

Ringtail possums appear highly susceptible to infection with *M. ulcerans* and are important reservoirs of the bacteria in Victoria. A One Health approach is needed to design and implement integrated interventions that reduce *M. ulcerans* transmission in Victoria, thereby protecting wildlife and humans from this emerging zoonotic disease.

**Author summary:** Buruli ulcer (BU), a neglected tropical skin disease, is emerging as a public health concern in the temperate Australian state of Victoria. Here, BU is spread by mosquitoes, and native possums are wildlife reservoirs of the causative bacterium, *Mycobacterium ulcerans*. Possums can be infected by BU, but knowledge of infection and disease in these animals is limited. We conducted surveillance studies in the two largest cities of Victoria, examining live and deceased possums. We found skin lesions in a third of examined possums and confirmed presence of *M. ulcerans* in almost three-quarters of the animals. Mouth swabs were positive for the bacteria in nearly two thirds of possums, as were pouch swabs of almost half the females. We also conducted mapping of the bodily distribution of skin lesions and found that paws and the undersides of abdomens and tails were the most affected areas. Our findings add support to the concept that possums, particularly ringtails, are *M. ulcerans* reservoirs in Victoria, and suggest several possible routes of infection for free-living possums that warrant further research. Improved understanding of BU in possums may allow development of targeted interventions that reduce disease transmission and protect both animal and human health.

## Introduction

*Mycobacterium ulcerans* is the causative agent of Buruli ulcer (BU), a progressive necrotising disease of skin and subcutaneous tissue and occasionally bone[1]. *M. ulcerans* is an environmental pathogen that can be introduced into a host’s subcutaneous tissues via puncture wounds, lacerations or other intradermal inoculating events such as insect bites[1, 2], although exact source(s) of infection and mode(s) of transmission have not yet been fully elucidated[3]. BU usually begins as a small skin nodule or swelling, first appearing several months after infection[1, 4]. Ulcerations can be extensive but are often painless, due to the locally cytotoxic, anaesthetic and immunosuppressive effects of the mycolactone toxins produced by *M. ulcerans* [5–7]. Without effective treatment, BU can lead to significant scarring and functional deformities, especially when involving bones or joints[1, 8].

Most human BU cases are reported in rural areas of western and central Africa [9–11], and in peri-urban areas of Australia, which is the only high-income country to report significant ongoing transmission[10]. In parts of Australia, BU is also known locally as Bairnsdale, Mossman or Daintree ulcer. Areas reporting limited *M. ulcerans* transmission have historically included the northern tropical regions of Queensland, Western Australia and the Northern Territory[12–14], with the first cases of autochthonous transmission retrospectively confirmed in 2021 in Batemans Bay, on the southern coast of New South Wales[15]. Since the early 2000s, the temperate south-eastern state of Victoria has been the country’s major BU hotspot, with the number, severity and geographical distribution of human cases increasing significantly in the past decade[16, 17]. Cases are now frequently reported in the state’s capital, Melbourne, and second largest city Geelong, and Victoria’s 2023 caseload was the highest on record with 362 cases, surpassing the previous peak of 340 cases in 2018[16].

BU is not restricted to humans. Small numbers of clinical BU cases have been reported in dogs, a cat, horses, alpacas, koalas and a potoroo[18–23] in Victoria, Australia. *M. ulcerans* DNA has been detected by PCR and genotype sequencing in lesion samples from mice in Ghana and Côte d’Ivoire[24, 25], and a dog and a goat in Benin[26]. Numerous studies from western and central Africa that tested hundreds of plant, invertebrate and animal samples detected *M. ulcerans* DNA only in small numbers of predatory aquatic insects, fish, and faeces from small rodents including grasscutters (*Thryonomys swinderianus*)[27–32]. These data suggest that the role of non-human animals in BU transmission in African settings may be limited.

In contrast, recent studies have demonstrated that BU is a vector-borne zoonosis in Victoria, Australia, with mosquitoes and native possums as bacterial vectors and reservoirs, respectively [19, 33–37]. While exact mechanisms of *M. ulcerans* transmission remain unclear, spatial clustering analysis has shown a striking overlap between clusters of human BU cases, *M. ulcerans*-harbouring mosquitoes and *M. ulcerans*-positive possum faeces in BU endemic areas of Victoria[33, 37]. *Aedes notoscriptus*, the ‘Australian backyard mosquito’, is capable of inoculating *M. ulcerans* through superficially contaminated skin during blood-feeding[38] and is the most consistently *M. ulcerans*-positive mosquito species in the Victorian BU-endemic area, at a reported frequency of up to 1%[33]. Common ringtail (CRT, *Pseudocheirus peregrinus*) and to a lesser extent common brushtail (CBT, *Trichosurus vulpecula*) possums, both of which are endemic and abundant in urban areas of Victoria, excrete large quantities of *M. ulcerans* in their faeces[19, 34], thus providing a potential source of environmental contamination for transmission to humans. Furthermore, the presence of *M. ulcerans*-positive possum excreta in a geographical area may be predictive of future human BU cases[37] possum excreta positivity in historically negative areas can predate the emergence of human BU cases by up to 31 months [39]. Faecal carriage of *M. ulcerans* in possums may be transient and can occur without accompanying clinical disease[36, 40]. Clinical BU in possums can manifest as mild, single, small shallow ulcers on a non-critical body site such as tail tip; to severe, multiple deep ulcerations on critical sites such as faces and distal limbs that can involve bones and joints and inhibit the climbing and foraging abilities of these arboreal animals[41]. CRTs may be more susceptible to infection and disease[36, 40] than other possum species. Systemic distribution of *M. ulcerans* throughout major internal organs has also been reported in CRTs, which may further reduce animal fitness and welfare[36, 41].

Despite recent advances in our understanding of *M. ulcerans* transmission in Victoria, knowledge of infection and disease in possums remains limited. This report expands on the previous research in this field by describing the major findings from surveillance studies conducted on free-living possums from endemic peri-urban areas of Victoria, south-eastern Australia.

## Materials and methods

Opportunistic necropsy-based and targeted trap-and-release surveillance studies were conducted on free-living possums in Victoria, Australia, between November 2021 and December 2022. This work was conducted as part of the ‘Beating Buruli in Victoria’ project led by the University of Melbourne’s Peter Doherty Institute for Infection and Immunity (Doherty Institute).

### Categorisation of possum BU status

Possums that had ulcerative skin lesions and for which at least one swab or tissue sample tested PCR positive for *M. ulcerans* DNA were designated as ‘clinical BU’. Possums without cutaneous lesions, but with at least one PCR positive *M. ulcerans* swab or tissue were designated as ‘infected’. Possums without positive PCR results from any swab or tissue sample were designated as ‘uninfected’. ‘PCR positive’ possums had at least one PCR positive swab or tissue sample, thereby encompassing both ‘clinical BU’ and ‘infected’ possum categories.

### Sourcing of cadavers

Cadavers of possums of any species that died or were euthanised for any reason at the U-Vet clinic in Werribee and the Lort Smith Animal Hospital in North Melbourne, Victoria, between November 2021 and December 2022 were collected and stored at -20°C. Recorded details included the possum’s geographical location, date and method of death, and clinical history where available. Cadavers were transported frozen to the Melbourne Veterinary School’s Anatomic Pathology lab and stored at -20°C.

### Live trapping events

Two possum trapping events were conducted during this study: the first was at the Barwon Heads Caravan Park in March 2022, a previously known focus of BU in humans, animals and mosquitoes[16, 19, 36, 42, 43], and the second took place in December 2022 on a residential property in Essendon where several previous BU-positive CRTs had originated[41, 44]. Trapping and sampling were conducted as per previously published methods[19]. Briefly, possum species-specific wire cages (smaller for CRTs, secured in trees or along fences; larger for CBTs, placed on the ground) were set at dusk and baited with apple pieces smeared with peanut butter. All cages were examined prior to dawn the next morning, and trapped possums were transferred to individual fabric bags and brought to the examination area. Voided faecal pellets were collected from traps or holding bags and placed into individually labelled ziplock bags. Each possum was placed under gaseous anaesthesia (isoflurane in oxygen, administered via a face mask) and subjected to a general clinical examination. Information was recorded on customised trapping examination sheets (S1 Appendix) including species, sex, weight, estimation of five-point body condition score (BCS)[45], and pouch check where applicable. Animals were classified as one of three age categories, based on body weight [46]: adult, sub-adult (fully weaned and independent of mother but not yet sexually mature), or juvenile. When present, back young were visually examined, weighed and an oral swab was collected (S1 Appendix); pouch young were not examined, and care was taken not to disturb them during the collection of the mother’s pouch swab.

Adult and sub-adult possums were microchipped subcutaneously between the scapulae, and plain microbiological swabs were collected from the oral cavity, cloaca, pouch where applicable, and any cutaneous lesions suggestive of BU. Where present, lesions were photographed, measured and described on the mapping silhouette on the examination record sheet, and a semi-subjective estimate of the severity of the lesions and associated welfare impacts recorded as mild, moderate or severe (see **Table 1**). These categories were adapted from the World Health Organization’s BU disease classification for assessment of human cases[47].

**Table 1:** Categorisation of the severity of ulcerative skin lesions in possums examined during this study.

| Category | Description |
| --- | --- |
| Mild | Single or multiple small lesions restricted to non-critical body sites (e.g. tail) |
| Moderate | Larger single or multiple small lesions that involve a critical site(s) (e.g. face, genitalia, footpad) but are unlikely to impair natural behaviours (e.g. climbing, foraging, feeding) |
| Severe | Very large single or multiple lesions at critical sites, and any lesions exposing bones or joints, that are likely to impair natural behaviours |

Healthy and mild- to moderately-afflicted possums were eligible for release after dusk at their trapping location. Where deemed necessary, euthanasia of severely compromised possums was conducted via intravenous or intracardiac injection of sodium pentobarbitone (Lethabarb®, Virbac Australia) while anaesthetised, and cadavers were transported to the Melbourne Veterinary School and stored at -20°C for inclusion in the necropsy study.

### Possum necropsies

The day prior to necropsy, cadavers were placed in a biosecurity cabinet and allowed to thaw at room temperature. All necropsies were conducted by the same veterinarian (ECH). A general external examination was conducted initially, and details of species, sex, weight, age categorisation (adult, sub-adult or juvenile, based on body weight[45]), estimation of five-point BCS[45], and pouch check where applicable were recorded on customised necropsy record sheets (S1 Appendix).

Where present, cutaneous lesions were first photographed, measured and described on the mapping silhouette on the necropsy record sheet, and a semi-subjective estimate of the severity of the lesions and associated welfare impacts recorded as mild, moderate or severe (see **Table 1**). Lesions were then swabbed, and multiple sections dissected and placed into formalin for histopathology, and into cryovials and stored at - 80°C prior to PCR testing. New scalpel blades were used prior to commencing each sampling step during necropsy.

Routine sample collection included plain microbiological swabs of each possum’s oral cavity, cloaca and pouch (where applicable), and dissection of a duplicate set of tissue and organ sections (full list in S1 Appendix) placed into formalin for histopathology, and into cryovials stored at -80°C prior to PCR testing, and culture where applicable. Samples of thoracic body cavity fluid, gut contents and faeces collected from the distal rectum were also collected into cryovials and stored at -80°C prior to PCR testing.

### Laboratory testing methods and definitions

#### DNA extraction and IS2404 PCR testing

All microbiological swabs, faecal pellets and frozen necropsy samples were transported on ice to the Mycobacterium Reference Laboratory at the Doherty Institute, Melbourne, Australia. DNA was extracted from these samples using a DNeasy® Blood and Tissue kit (Qiagen, Germany) or DNeasy® PowerSoil kit (Qiagen, Germany). Procedural extraction control blanks (sterile water) were included to monitor potential PCR contamination. The extracted DNA samples were screened by real-time PCR targeting the IS2404 insertion sequence in the *M. ulcerans* genome, the standard assay for molecular detection of *M. ulcerans*, using methods as described[48].

Most samples were tested in duplicate, and results were categorised as ‘positive’ when the average cycle threshold (Ct) of both duplicates was ≤40[48]; ‘negative’ when *M. ulcerans* was not detected or the average Ct of both duplicates was >40; and ‘equivocal’ when one duplicate was positive and the other negative. When only single samples were tested, results were either ‘positive’ when the Ct was ≤40, or ‘negative’ for Cts >40.

#### Culture of PCR positive tissues

A subset of PCR positive samples was subjected to culture. Portions of IS2404 positive tissue samples were placed in screw cap tubes containing 0.5g glass beads and 600ul 1x phosphate buffered saline. Tubes were subjected to 4 rounds of homogenisation in a Precellys 24 bead beater (Bertin Technologies, France) for 30 seconds at 6500rpm each. 300ul of this homogenate was transferred to a fresh tube containing 300ul of 2% NaOH and incubated for 15 minutes at room temperature. Drops of 10% orthophosphoric acid was added to neutralise the solution. 50-100μl aliquots of the treated samples were then plated out on to Middlebrook 7H10 agar with PANTA and Brown and Buckle slopes, incubated at 30°C for 8-10 weeks and then any observed bacterial growth was sub-cultured, genomic DNA extracted and whole gene sequencing performed.

#### Immunohistochemical staining and histological examination

Formalin-fixed tissues were routinely processed and stained by haematoxylin-eosin, Gram and Ziehl-Neelsen stains and examined histologically by a veterinary pathologist.

### Statistical analysis

The data from both the possum trapping and necropsy studies were combined, and the categorical variables of a) possum species (CRT and CBT), b) sex (male and female), c) age category (adult, sub-adult, juvenile) and d) BU status (‘clinical BU’, ‘infected’ and ‘uninfected’) were compared using 2x2 contingency tables. The two-tailed Fisher’s exact test was used to test for associations between categorical variables in inter-group comparisons. The P value was set at ≤0.05.

### Ethics statement

Collection and examination of deceased possums was conducted under DEECA permit numbers 10009447 and 10010578. Prior to the commencement of possum trapping, ethics approval was granted by the University of Melbourne Faculty of Science’s animal ethics committee LAE2 (project ID 22910), and permission to conduct trapping and sampling of free-living possums was obtained from DEECA (wildlife research permit number 10010257).

## Results

### Possum trapping study

#### Trapping site 1: Barwon Heads

Ten CRT traps and five CBT traps were set around the Barwon Heads Caravan Park overnight on 30 March 2022, and five possums were trapped: three adults, including one with a back joey, and a sub-adult (see **Table 2**.) Four of the possums were CBTs and one, an adult, was a CRT. Most were in fair to good condition, except for the sub-adult male CBT (T3) which was very thin (BCS 1.5/5) and had a heavy flea infestation around the ears, eyes and muzzle.

**Table 2:**
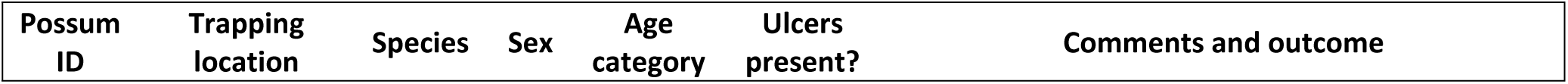
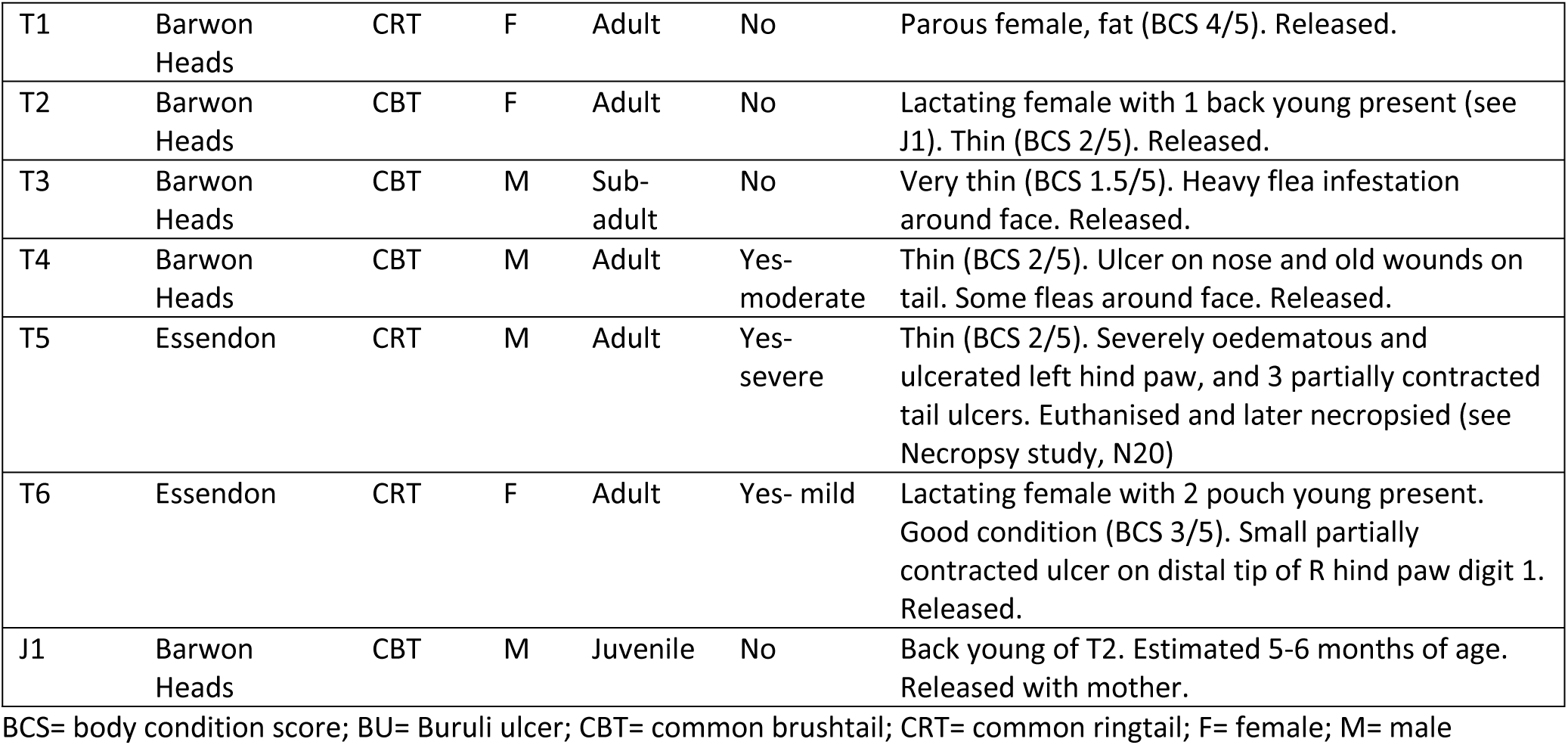
Descriptive data for possums examined during the two live trapping studies, Victoria, 2022.

Cutaneous lesions suggestive of BU were only observed on the adult male CBT (T4): the nasal rostrum bore a full-thickness ulcerative lesion measuring approximately 5mm long and 3mm wide, and the ventral mid-section of the tail bore several dry, partially scabbed wounds. The ulcerative lesions were assessed to be of moderate severity and to have similar welfare impacts, not necessitating euthanasia. All five possums trapped in this location were subsequently released.

The faecal pellet from the adult male CBT, T4, was positive for *M. ulcerans* by IS2404 PCR, resulting in this possum being categorised as ’clinical BU’. All other samples from this possum, including lesion and cloacal swabs, were negative. All other faecal pellets and swabs collected from the remaining possums trapped at Barwon Heads tested negative for *M. ulcerans* by IS2404 PCR, and these four possums were therefore classified as ‘uninfected’ (see Table 3). Fleas taken from the facial regions of T3 and T4 were identified as stick-tight fleas (*Echidnophaga gallinacea*).

**Table 3:** IS2404 PCR results and BU status for possums examined during the two live trapping studies, Victoria, 2022.

| Possum ID | Ct results<sup>a</sup> (average<sup>b</sup>) |  |  |  |  |  | BU status |
| --- | --- | --- | --- | --- | --- | --- | --- |
|  | Voided faeces | Oral swab | Cloacal swab | Pouch swab | Ulcer swab | Ulcer location |  |
| T1 | Negative | Negative | Negative | Negative | N/A |  | Uninfected |
| T2 | Negative | Negative | Negative | Negative | N/A |  | Uninfected |
| T3 | Negative | Negative | Negative | N/A | N/A |  | Uninfected |
| T4 | 34.04 | Negative | Negative | N/A | Negative |  | Clinical BU |
| T5 | 22.22 | 33.85 | 35.12 | N/A | 23.83<br>29.28 | Hind paw<br>Tail | Clinical BU |
| T6 | 26.12 | Equivocal | 38.85 | Negative | 38.80 | Hind paw | Clinical BU |
| J1 | N/A | Negative | N/A | N/A | N/A |  | Uninfected |

Ct= real-time PCR cycle threshold; N/A= not applicable.
<sup>a</sup>Negative= Ct≥40; Equivocal= one positive and one negative PCR result.
<sup>b</sup>Average of two tested duplicates.

#### Trapping site 2: Essendon

Eight CRT traps were set around the Essendon residential property overnight on 13 December 2022, and two possums were trapped: both were adult CRTs and both bore cutaneous lesions suggestive of BU (see Table 2). The male (T5) was in poor body condition and had a severely oedematous left hind paw; palpation indicated disruption of the internal structure, and purulent material exuded from the linear ulcerative sinus present on the dorsal aspect. The severity of this lesion necessitated euthanasia, and the cadaver was transported to the Melbourne Veterinary School at Werribee for necropsy (necropsy case number N20; see following section). Three small, shallow and partially contracted lesions were also observed on the lateroventral tail.

The second CRT trapped in this location (T6) was a young lactating female in good body condition, with two pouch young. A small, partially contracted shallow ulcer was observed on the distal tip of the first digit (‘thumb’) of the right hind paw. Given the mild lesion severity and associated welfare impacts, this possum was later released.

All samples from T5 were PCR positive for *M. ulcerans*, with notably low Ct values recorded for the hind paw ulcer swab and voided faecal pellet, indicative of a high bacterial load. While T6’s pouch swab was negative and her oral swab was equivocal, her faecal pellet, cloacal swab and the swab taken from her hind paw ulcer were all PCR positive. Both T5 and T6 were therefore categorised as ‘clinical BU’ (see Table 3).

Seven mosquitoes opportunistically collected during the Essendon trapping event were also tested for the presence of *M. ulcerans* DNA. Two returned positive results (average Cts 33.9 and 37.0), and another was equivocal (S2.1 table).

### Necropsy study

A total of 20 possums were necropsied, including the male CRT that was trapped and euthanised during the Essendon trapping event (T5/N20). Eighteen possums (90%) were CRTs, and most (n=16, 80%) originated from inner suburbs of Melbourne (Fig. 1). Equal numbers of male and female possums were examined. Four (20%) were adults, ten (50%) were subadults, and six (30%) were juveniles. Full or partial histories were available for all but two possums (see Table 4).

**Fig. 1:**
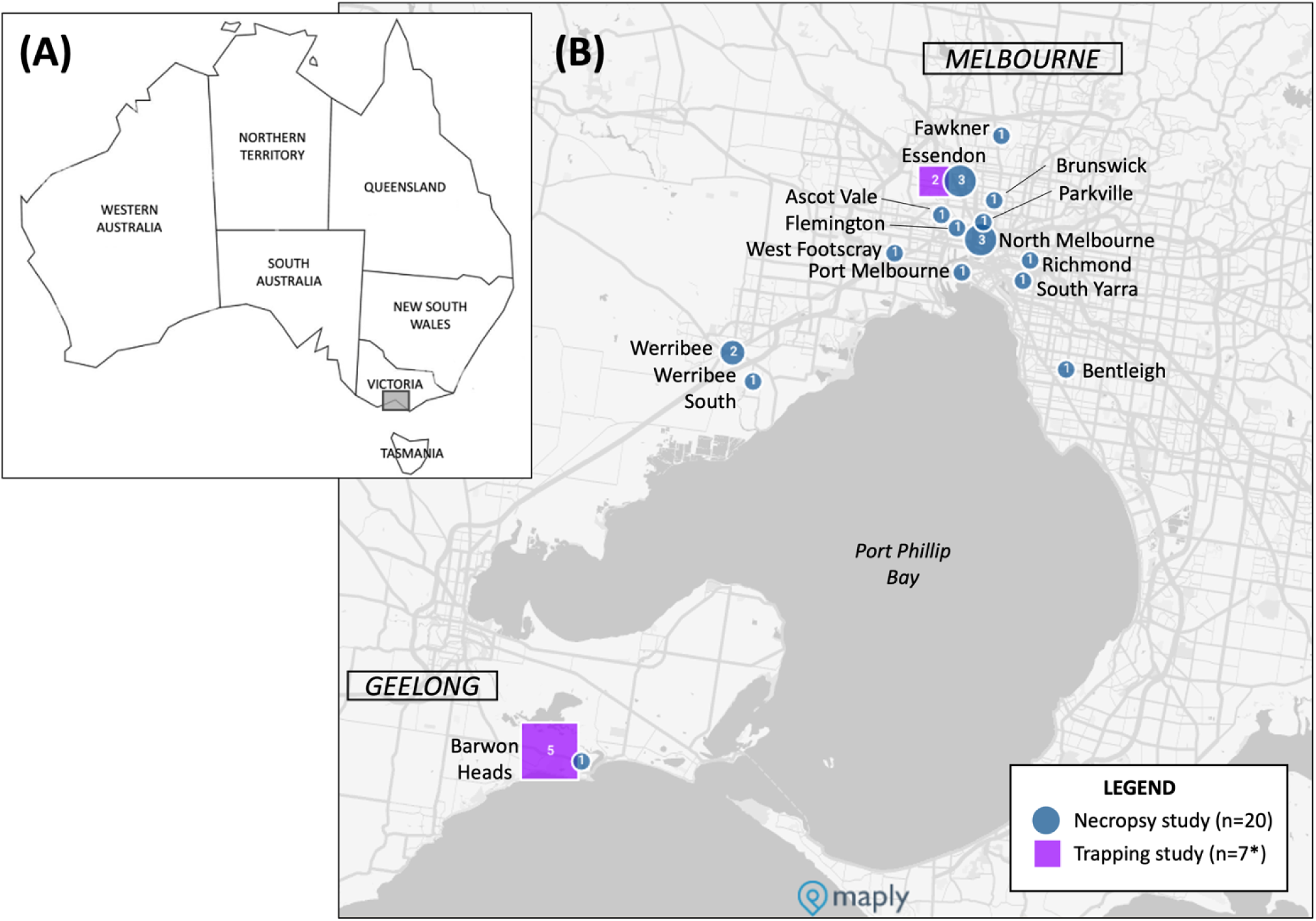
Maps showing the locations from which the possums examined in this study (2021-2022) originated. (A) Map of Australia, with the study area within the south-eastern state of Victoria highlighted by the grey box. (B) Inset shows the region of Melbourne and Geelong from which the possums examined in this study originated. Possums examined during the live trapping are shown as purple squares while the those included in the necropsy study are shown as blue circles. The size of the icons reflect increasing numbers of possums from that location and the numbers inside the icons state the number of possums from each location. *One possum was trapped in Essendon, then euthanized and included in the necropsy study, so is counted twice in this figure.

**Table 4:** Descriptive data for possums examined during the necropsy study, 2021-2022.

| Case ID | Suburb of origin | Species | Sex | Age category | Ulcers present? | History and necropsy findings |
| --- | --- | --- | --- | --- | --- | --- |
| N1<sup>a</sup> | Essendon | CRT | M | Adult | Yes – severe | Captured and euthanised, due to deep ulcers on scrotum and R hind paw, with tarsal bone of digit 1 (‘thumb’) exposed. Presumptive diffuse pulmonary microgranulomas. Thin (BCS 2/5). |
| N2 | Werribee | CRT | F | Adult | No | Found dead. Thin (BCS 2/5); 3 pouch young present. Trauma and S/C haemorrhage around head, neck and thorax. |
| N3 | Barwon Heads | CBT | F | Juvenile | No | Found dead. Back joey, estimated 6mo. Puncture wound and S/C haemorrhage into R lateral thorax. |
| N4 | Werribee | CRT | F | Subadult | No | Died in care. Fair condition (BCS 2.5/5); tapeworm in proximal small intestine. |
| N5 | Werribee South | CRT | M | Subadult | No | No history available. Very thin (BCS 1.5/5); abdominal trauma and S/C haemorrhage. |
| N6<sup>a</sup> | Bentleigh | CRT | M | Adult | Yes – severe | Captured and euthanised due to multiple deep ulcers: L fore paw with carpal bones exposed, R fore paw, L and R hind paw, scrotum, tail. Thin (BCS 2/5). |
| N7 | South Yarra | CRT | F | Subadult | No | Found crushed by pallet, euthanised. Thin (BCS 2/5). Multiple open tail fractures, S/C haemorrhage around abdomen and pelvis. |
| N8 | Richmond | CRT | F | Adult | Yes – moderate | Found on ground, euthanised due to swelling and ulceration to nasal dorsum, ulcer on distal tail. Fair BCS (2.5/5). |
| N9 | Parkville | CBT | M | Juvenile | No | Found on ground, cold and unresponsive, euthanised. Estimated 4.5mo, emaciated (BCS 1), fractured skull and cranial haemorrhage. |
| N10 | Ascot Vale | CRT | F | Adult | Yes – mild | Found on ground with neurological defects, euthanised. Very thin (BCS 1.5/2), evidence of head trauma. Two small shallow ulcers on distal tail tip. |
| N11 | North Melbourne | CRT | M | Juvenile | Yes – mild | Found injured on ground, euthanised. Back joey, estimated 5mo. Bite marks and S/C haemorrhage around head. Small shallow ulcer on distal tail tip. |
| N12 | Port Melbourne | CRT | M | Juvenile | No | Euthanised but no history available. Back joey, estimated 5mo, thin (BCS 2/5). Extensive cranial trauma: multiple skull fractures, brain haemorrhage, R eye protruding from socket. |
| N13 | Fawkner | CRT | F | Subadult | No | Found on park bench with traumatic amputation of tail and myiasis, euthanised. Fair condition (BCS 2.5/5). Lungs very firm, white froth throughout. Tapeworms (n=2) in distended proximal small intestine. |
| N14<sup>a</sup> | Essendon | CRT | F | Adult | Yes – severe | Found on road being attacked by birds, euthanised due to multiple deep ulcerations: R fore paw, L and R hind paws, distal tail. Emaciated (BCS 1/5). Patchy pulmonary congestion and oedema. |
| N15 | Flemington | CRT | M | Juvenile | No | Found on ground, died in care. Back joey, estimated 4.5mo, fair condition (BCS 2.5/5). |
| N16 | North Melbourne | CRT | M | Adult | No | Found on ground with traumatic amputation of tail and myiasis, euthanised. Thin (BCS 2/5). |
| N17 | West Footscray | CRT | F | Adult | No | Found on ground with pelvic trauma, euthanised. Parous female, good condition (BCS 3/5). S/C haemorrhage and fractures to abdomen and pelvis. |
| N18 | Brunswick | CRT | F | Juvenile | No | Found on ground, condition deteriorated in care, euthanised. Back joey, estimated 7mo, distended abdomen and digestive tract. |
| N19 | North Melbourne | CRT | M | Adult | No | Found on ground with injured back leg, euthanised. Good condition (BCS 3/5). S/C haemorrhage to ventrum. |
| N20<sup>a</sup> (T5) | Essendon | CRT | M | Adult | Yes – severe | Captured during Essendon trapping event (T5), euthanised. Severely oedematous L hind paw, exuding purulent material from linear ulcerative opening on dorsal surface that extended to a central cavity exposing ligaments and tendons on dissection. Three partially contracted ulcers on lateroventral surface of proximal tail. Thin (BCS 2/5). |

BCS= body condition score; BU= Buruli ulcer; CBT= common brushtail; CRT= common ringtail; F= female; M= male; mo= months of age; S/C= subcutaneous; L= left; R= right
<sup>a</sup>Case described in more detail, including photographs of ulcerative lesions, in [41].

#### Necropsy findings and lesion mapping

Two thirds (n=13, 65%) of the necropsied possums showed no lesions suggestive of BU and had died or were euthanised for other reasons, predominantly traumatic injuries. Tapeworms present in the proximal small intestine of two cases (N4, N13, both subadult female CRTs) were identified as *Bertiella paraberrata*[49]. Diffuse white nodules, approximately 1mm in diameter, were observed grossly throughout the lungs of N1 and interpreted as presumptive microgranulomas[41].

Ulcerative lesions suggestive of BU were observed in 35% (7/20) of the necropsied possums. Lesions observed in four of these possums are shown in Fig. 2. Of the seven ulcerated possums, five were euthanised due to the severity of multiple deep ulcerative lesions and/or swelling diagnosed as presumptive BU by the examining veterinarians: four severe cases with multiple, deep severe cutaneous ulcerations to one or more paws (N1, N6, N14 and N20), which in two cases (N1, N6) exposed carpal and tarsal bones; and one moderate case (N8) that had swelling and ulceration of the nasal dorsum, and a small tail ulcer. Two mild cases (N10 and N11) had small ulcers on the distal tail tip which were incidental findings; both possums had been euthanised due to unrelated traumatic injuries.

**Fig. 2:**
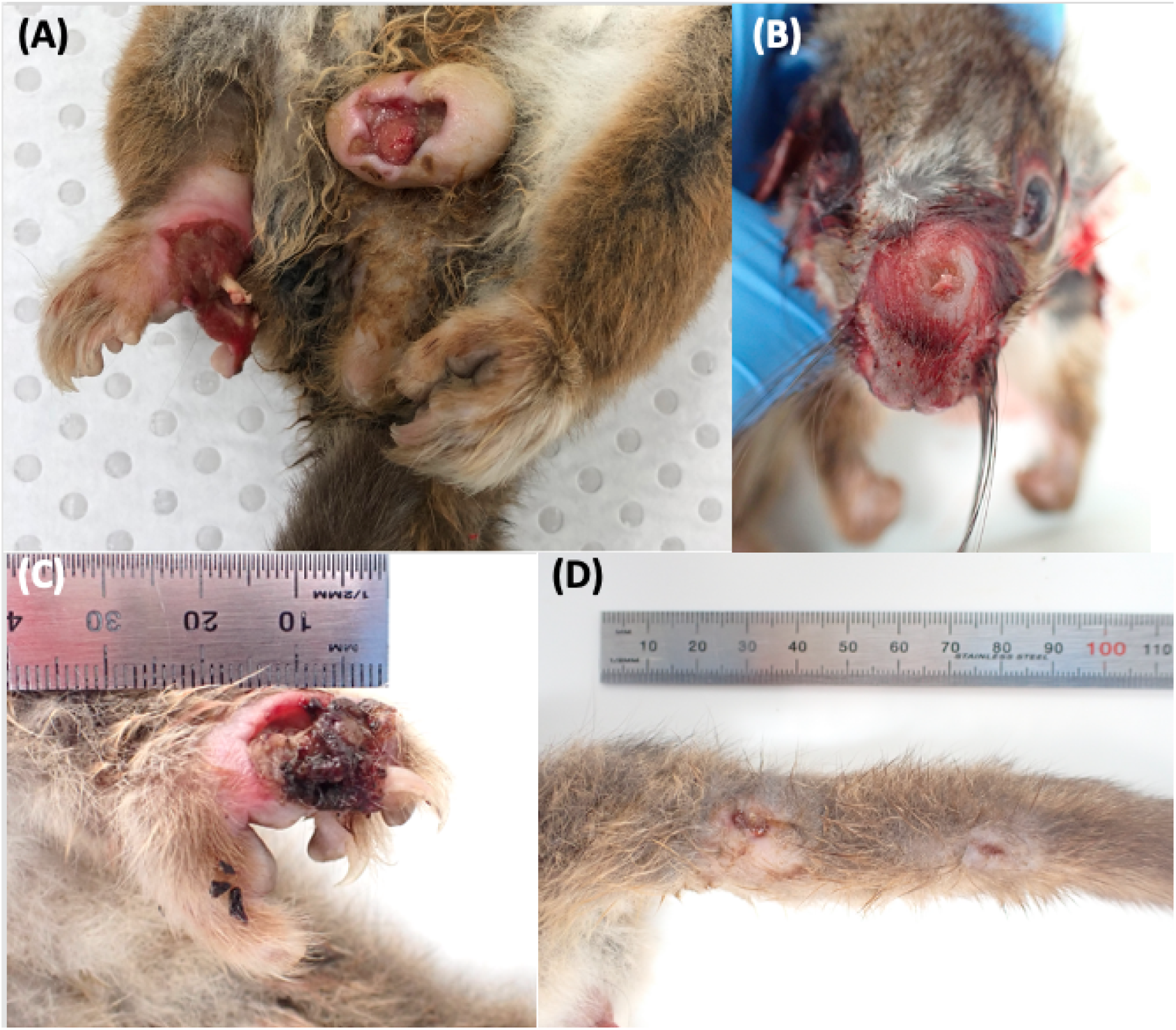
Buruli ulcer lesions in four of the necropsied common ringtail possums. (A) Scrotal and right hind paw ulcers (N1). The first metatarsal bone of the right hind paw is exposed and only a skin flap remains of the first digit (See [41]). (B) Moderate swelling and ulceration of the dorsal nasal rostrum (N8). (C) Deep ulceration of the dorsal aspect of the left fore paw (N14), with deeply undermined caudal wound edges. A plug of necrotic material is partially covering exposed striated muscle and tendons. (D) Two partially healed ulcers on the proximal lateroventral aspect of the tail (N20/T5).

Overall, 24 lesions were observed on the 7 clinically affected possums. Lesion mapping revealed that distal limbs and tails were the most frequently affected regions (see Fig. 3). Nearly half of all observed lesions (11/24, 46%) were on the distal limbs, and all were either moderate or severe: three on the carpi, of which one exposed carpal bones and another exposed striated muscle and tendons; and eight on the tarsi, of which one exposed tarsal bones. Tail lesions (10/24, 38%) were all located on the ventral or ventrolateral aspect, and all were mild. The remaining three ulcers were located on the nasal dorsum, cloaca, and scrotum. All but one of the clinically affected possums had two or more ulcers; the one possums with a single ulcers had a small lesion on the distal tail tip.

**Fig. 3:**
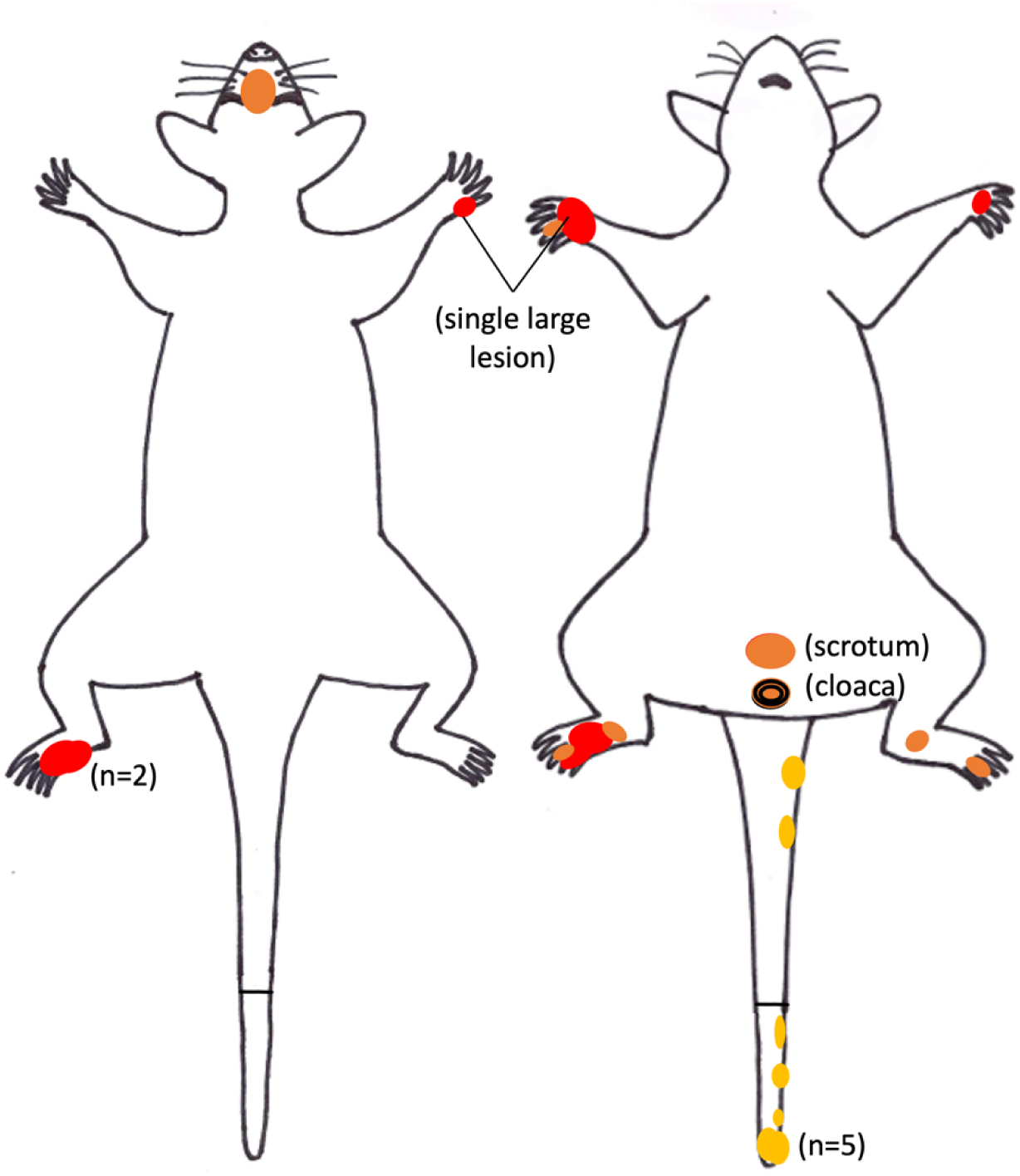
Possum mapping silhouette showing cutaneous ulcerative lesions recorded in the seven clinically affected possums examined during the necropsy study, 2021-2022. The coloured circles depict the approximate size and location of the ulcerative lesions. The colour of the circle denotes severity of lesions: yellow for mild/shallow; orange for moderate; red for severe/deep. Where more than one circle occurs in the same location, the number of overlapping circles (n) is stated.

#### IS2404 PCR testing of swabs

A total of 59 swabs were collected from the 20 necropsied possums: from oral cavities (n=20), cloacas (n=20), pouches (n=9), and ulcerative lesions (n=10, from the 7 clinically affected possums). Five possums, all of which had severe or moderate clinical BU, returned positive PCR results for every swab that was collected (see Table 5).

**Table 5:**
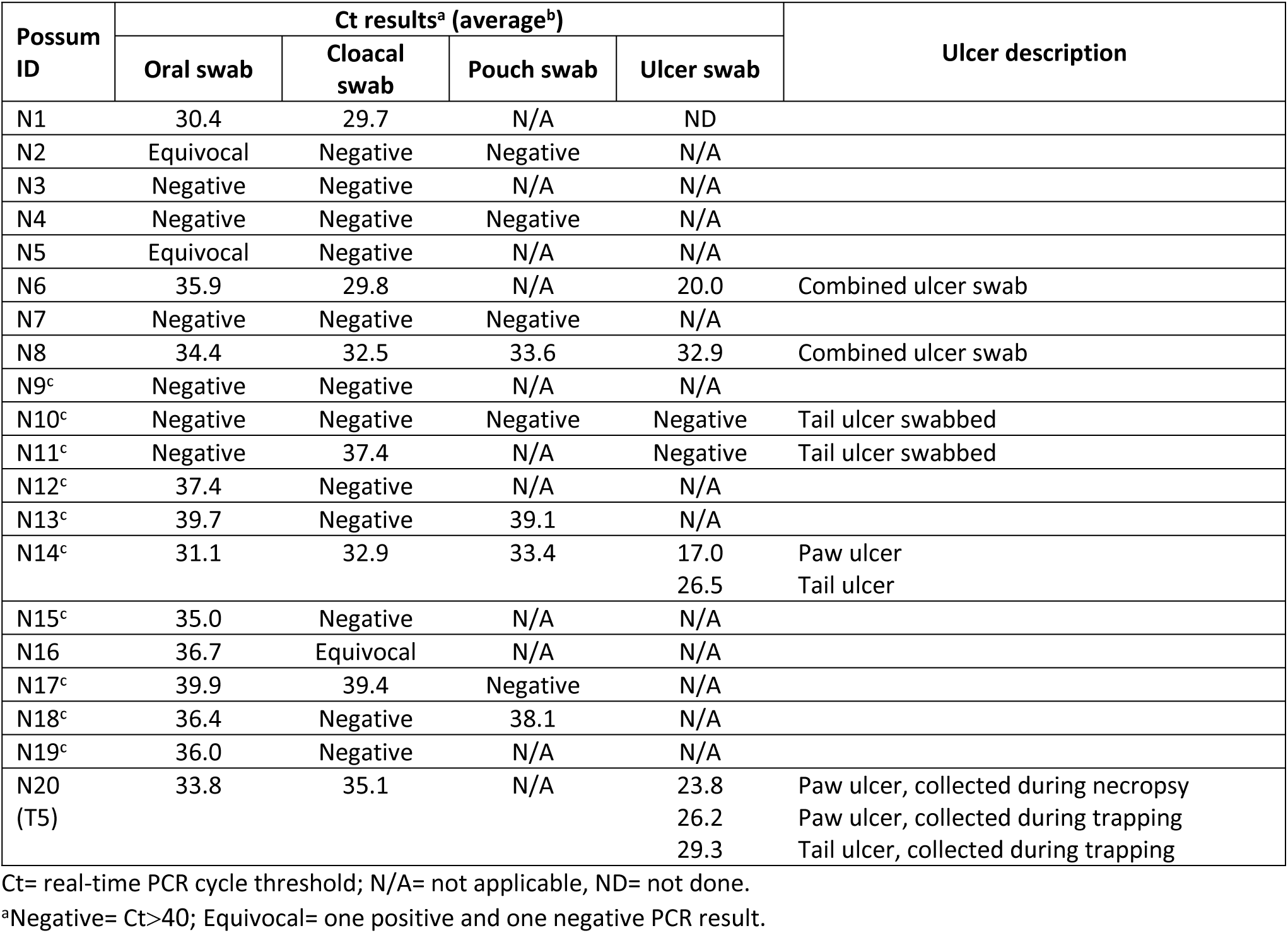
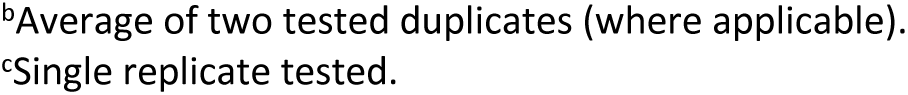
IS2404 PCR results from swabs collected during the possum necropsy study.

| Possum ID | Ct results<sup>a</sup> (average<sup>b</sup>) |  |  |  | Ulcer description |
| --- | --- | --- | --- | --- | --- |
|  | Oral swab | Cloacal swab | Pouch swab | Ulcer swab |  |
| N1 | 30.4 | 29.7 | N/A | ND |  |
| N2 | Equivocal | Negative | Negative | N/A |  |
| N3 | Negative | Negative | N/A | N/A |  |
| N4 | Negative | Negative | Negative | N/A |  |
| N5 | Equivocal | Negative | N/A | N/A |  |
| N6 | 35.9 | 29.8 | N/A | 20.0 | Combined ulcer swab |
| N7 | Negative | Negative | Negative | N/A |  |
| N8 | 34.4 | 32.5 | 33.6 | 32.9 | Combined ulcer swab |
| N9<sup>c</sup> | Negative | Negative | N/A | N/A |  |
| N10<sup>c</sup> | Negative | Negative | Negative | Negative | Tail ulcer swabbed |
| N11<sup>c</sup> | Negative | 37.4 | N/A | Negative | Tail ulcer swabbed |
| N12<sup>c</sup> | 37.4 | Negative | N/A | N/A |  |
| N13<sup>c</sup> | 39.7 | Negative | 39.1 | N/A |  |
| N14<sup>c</sup> | 31.1 | 32.9 | 33.4 | 17.0<br>26.5 | Paw ulcer<br>Tail ulcer |
| N15<sup>c</sup> | 35.0 | Negative | N/A | N/A |  |
| N16 | 36.7 | Equivocal | N/A | N/A |  |
| N17<sup>c</sup> | 39.9 | 39.4 | Negative | N/A |  |
| N18<sup>c</sup> | 36.4 | Negative | 38.1 | N/A |  |
| N19<sup>c</sup> | 36.0 | Negative | N/A | N/A |  |
| N20 (T5) | 33.8 | 35.1 | N/A | 23.8<br>26.2<br>29.3 | Paw ulcer, collected during necropsy<br>Paw ulcer, collected during trapping<br>Tail ulcer, collected during trapping |

Ct= real-time PCR cycle threshold; N/A= not applicable, ND= not done.
<sup>a</sup>Negative= Ct>40; Equivocal= one positive and one negative PCR result.
<sup>b</sup>Average of two tested duplicates (where applicable).
<sup>c</sup>Single replicate tested.

Of the standard swab types collected, the oral cavity swabs were the most commonly PCR positive (12/20, 60%), followed by the pouch swabs (4/9, 44%) (see S2.2 Table). Of the swabs taken from ulcerative lesions, 80% were positive, and this swab type had the lowest recorded Ct value (17.0, from N14 paw ulcer) of all swabs tested. The two negative ulcer swabs came from distal tail ulcers from the two mildly affected possums (N10, N11).

#### IS2404 PCR testing of tissues

A full suite of up to 20 tissue and organ samples were collected from most of the 20 necropsied possums and subjected to IS2404 PCR; results are presented in Table 6. In some cases, samples were not tested, either because a sample type was added to the collection protocol after the commencement of the necropsy study (e.g. heart), because samples were not collected or could not be found at the testing laboratory.

**Table 6:** IS2404 PCR results from tissue and organ samples collected during the possum necropsy study.

| Possum ID | Ct results<sup>a</sup> (average<sup>b</sup>) |  |  |  |  |  |  |  |  |  |  |  |  |  |  |  |  |  |  | No. positive tissues | % positive tissues |
| --- | --- | --- | --- | --- | --- | --- | --- | --- | --- | --- | --- | --- | --- | --- | --- | --- | --- | --- | --- | --- | --- |
|  | Ear | Nose | Front footpad | Hind footpad | Submand. LN | Heart | Lung | Liver | Spleen | Kidney | Adrenal gland | Body cav. fluid | Gut contents | Stomach | Small intestine | Caecum | Large intestine | Faeces | Ulcer(s) |  |  |
| N1 | 24.2 | 24.2 | 28.3 | 26.8 | 30.5 | ND | 31.3 | 32.5 | 34.6 | 33.8 | 30.1 | 38.7 | 21.6 | 25.2 | 30.6 | 28.0 | 29.1 | 13.5 | 15.0 | 18 | 100 |
| N2 | Neg | Neg | 39.6 | 37.1 | Neg | ND | Neg | 39.1 | 33.6 | Neg | Neg | ND | Eq | Neg | Neg | 33.7 | Neg | Neg | N/A | 5<sup>d</sup> | 31<sup>d</sup> |
| N3 | Neg | Neg | Neg | 40.0 | Neg | ND | Neg | Neg | Neg | Neg | Neg | Neg | Neg | 36.5 | Eq | Neg | Neg | Neg | N/A | 1<sup>d</sup> | 6<sup>d</sup> |
| N4 | Neg | Neg | Neg | Neg | Neg | ND | Neg | Neg | Eq | Neg | Neg | Neg | Neg | Neg | Neg | Neg | Neg | Neg | N/A | 0<sup>d</sup> | 0<sup>d</sup> |
| N5 | Neg | Neg | Neg | Neg | Neg | ND | Neg | 38.4 | Eq | Neg | Eq | Neg | 38.3 | Neg | Neg | 39.8 | 39.4 | Neg | N/A | 4<sup>d</sup> | 24<sup>d</sup> |
| N6 | 28.1 | 27.2 | 28.8 | 27.2 | 28.3 | 33.6 | 34.3 | 33.8 | 36.2 | 33.7 | 31.2 | 35.9 | 22.4 | 34.1 | 32.1 | 32.7 | 33.8 | 19.1 | 25.6 | 19 | 100 |
| N7 | Neg | Neg | Neg | Neg | Neg | Neg | Neg | Neg | ND | Neg | Neg | Neg | Neg | Neg | Neg | Neg | Neg | Neg | N/A | 0 | 0 |
| N8 | 32.0 | ND | 28.4 | 34.2 | 35.7 | Eq | 33.4 | 37.7 | Eq | 38.5 | 34.3 | Neg | Neg | 35.4 | 35.4 | Neg | 35.6 | 21.6 | ND | 12<sup>d</sup> | 71<sup>d</sup> |
| N9 | Neg | 37.9 | Neg | Neg | Neg | Neg | Neg | Neg | Neg | Neg | Neg | Neg | Neg | Neg | Neg | Neg | Neg | Neg | N/A | 1 | 6 |
| N10 | Neg | Neg | Neg | Neg | Neg | Neg | Neg | Neg | Neg | Neg | Neg | Neg | Neg | Neg | Neg | Neg | Neg | Neg | Neg | 0 | 0 |
| N11 | Neg | Neg | Neg | Neg | Neg | Neg | Neg | Neg | Neg | Neg | Neg | Neg | Neg | Neg | Neg | Neg | Eq | Neg | Neg | 0<sup>d</sup> | 0<sup>d</sup> |
| N12 | 38.8 | Neg | Neg | Neg | Neg | Neg | Neg | Neg | Neg | Neg | Neg | Neg | 38.7 | Neg | Neg | Neg | Neg | Neg | N/A | 2 | 11 |
| N13 | Neg | Neg | Neg | Neg | Neg | Neg | Neg | Neg | Neg | Neg | Neg | Neg | Neg | Neg | Neg | Neg | Neg | Neg | N/A | 0 | 0 |
| N14 | 24.2 | 25.8 | 28.1 | 26.4 | 27.8 | 29.9 | 26.4 | 30.6 | 31.8 | 34.6 | 33.1 | 30.3 | 22.6 | 32.3 | 34.4 | 32.3 | 31.9 | 19.5 | 19.5<sup>c</sup> | 19 | 100 |
| N15 | Neg | Neg | 39.2 | Neg | Neg | Neg | Neg | Neg | ND | Neg | Neg | Neg | Neg | Eq | Neg | Neg | Neg | Neg | N/A | 1<sup>d</sup> | 6<sup>d</sup> |
| N16 | Neg | Neg | Neg | Neg | Neg | Neg | Neg | Neg | Neg | Neg | Neg | Neg | Neg | Eq | Eq | Neg | Neg | Neg | N/A | 0<sup>d</sup> | 0<sup>d</sup> |
| N17 | Neg | Neg | Neg | Neg | Neg | Neg | Eq | Neg | Neg | Neg | Neg | Neg | Neg | Neg | Neg | Neg | Eq | Neg | N/A | 0<sup>d</sup> | 0<sup>d</sup> |
| N18 | Neg | Neg | Neg | Neg | Neg | Neg | Neg | Neg | Neg | Neg | Neg | Neg | Neg | Neg | Neg | Neg | Eq | Neg | N/A | 0<sup>d</sup> | 0<sup>d</sup> |
| N19 | Neg | Neg | Neg | Neg | Neg | Neg | Eq | Neg | Neg | Neg | Neg | 40.0 | Neg | Neg | Neg | Neg | Neg | Neg | N/A | 1<sup>d</sup> | 6<sup>d</sup> |
| N20 (T5) | 32.8 | 31.9 | 27.4 | 22.5 | 31.6 | 34.3 | 36.0 | 35.2 | 35.7 | 36.1 | 31.1 | 36.9 | 24.5 | 33.2 | 37.9 | 32.8 | 33.3 | 22.2 | 24.2<sup>c</sup> | 18 | 100 |

Body cav. fluid= body cavity fluid; Ct= real-time PCR cycle threshold; Eq= equivocal; N/A= not applicable; Neg= negative; ND= not done; Submand. LN= submandibular lymph node.
<sup>a</sup> Negative= Ct>40; Equivocal= one positive and one negative PCR result.
<sup>b</sup> Average of two tested duplicates (where applicable).
<sup>c</sup> Average of two sampled ulcers.
<sup>d</sup> Not including equivocal results.

The four clinically affected possums assessed as severe (N1, N6, N14, N20) all tested PCR positive in every tissue and organ sample that was tested, including faeces. One sample of bone marrow, collected from the right femur of possum N20, was also PCR positive (average Ct 36.98). The moderately affected possum (N8) was positive in 71% (12/17) of tested tissues; the exceptions were body cavity fluid, gut contents and caecum, which were negative, and heart and spleen, which were equivocal.

Seven possums with no cutaneous lesions had at least one tissue that tested PCR positive and were classified as ‘infected’: N2 had five (31%), N5 had four (24%), N12 had two (11%) and N3, N9, N15 and N19 each had one (6%) tissue that tested PCR positive. All but N9 and N12 also had at least one equivocal result. There was little consistency in the tissues and organs that tested positive for these possums: four sample types (front footpad, liver, gut contents, caecum) were positive for two of the possums, whereas the six remaining sample types (nose, hind footpad, spleen, body cavity fluid, stomach, large intestine) were each positive in only one possum.

The lowest Ct value, 13.5, was recorded from the faecal sample and testicular ulcer section collected from N1. This was the lowest recorded Ct value for any of the tested sample types (swabs, tissues and organ samples). Of the ten tissue and organ samples with the lowest recorded Ct values, five were faeces, two were ulcers, two were gut contents, and one was a section of skin from the hind footpad. Of the tissue and organ sections, dissected ulcers were the most frequently PCR positive (6/9, 66.7%) (see S2.2 Table).

#### Culture of PCR positive tissues

*M. ulcerans* was cultured from the testicular tissue of N1. Attempts to culture dissected lesion sections from N14 and N20 were unsuccessful.

#### Histopathological examination of tissue and organ samples

Examination of the dissected skin lesions from the four severe (N1, N6, N14, N20) and one moderate (N8) case confirmed the presence of moderate to exceedingly severe ulcerative necrotising pyogranulomatous dermatitis, panniculitis, cellulitis and myositis, with ulceration and inflammation expanding to the underlying bone in N6 and N8. Abundant Gram positive and acid-fast bacilli were observed on the surface and deep within the dermis and underlying tissues, concentrated in areas of tissue necrosis. Thick serocellular crusting was observed over the right hindlimb ulcer of N14 (see Fig. 4.) Sections taken from the left forelimb ulcer of N6 and the partially contracted proximal tail ulcers of N20 were observed to have mild epidermal hyperplasia of the ulcer margins, multifocal lymphoid aggregates in the subcutis and other changes consisted with ongoing cutaneous healing.

**Fig. 4:**
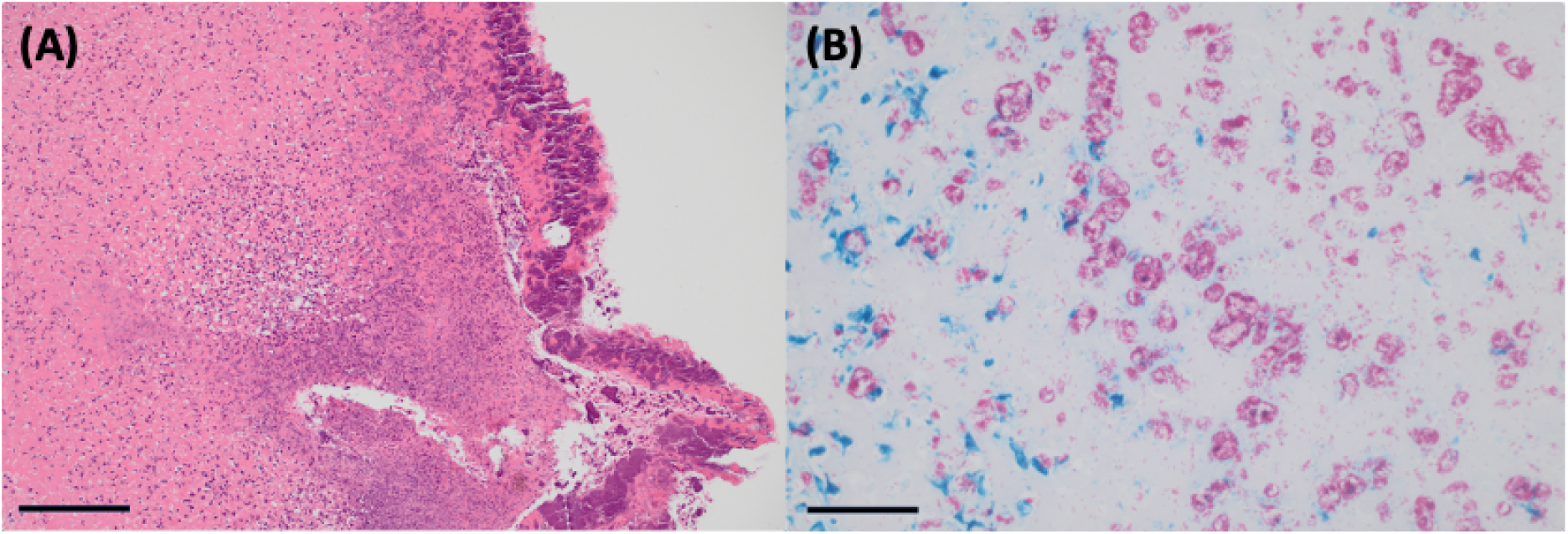
Stained sections of cutaneous lesions taken from a common ringtail possum with Buruli ulcer (N14). (A) An extensive, thick layer of serocellular crust covers and replaces the ulcerated epidermis. The crust consists of viable and degenerate mixed inflammatory cells, and cellular debris in which bacteria are enmeshed. The dermis is diffusely necrotic and infiltrated by variable numbers of mixed inflammatory cells and bacteria. Hematoxylin and eosin. Bar = 400 um. (B) Ziehl-Nielsen-stained sections of the sections shown in (A), highlighting the abundant presence of monomorphic acid-fast-positive bacteria (shown as pink structures) throughout the necrotic dermis. Ziehl-Nielsen. Bar = 200 um.

Widespread freeze-thaw artifact and autolysis were present to varying degrees in the tissues and organs of all examined possums, which prevented meaningful histological interpretation of many sections, including the lung tissue from N1. Nevertheless, evidence of systemic pathology was observed in four cases (see Table 7). The pancreas of N6 showed possible mild acinar atrophy, consistent with poor nutritional condition. N8 and N14 showed evidence of marked pulmonary oedema, with thrombosis of several medium calibre vessels and several mild lymphohistiocyctic aggregates also observed in N14, however granulomas were not definitively detected. Moderate neutrophilic, lymphohistiocytic hepatitis was also observed in N14, and multifocal, mild to moderate portal hepatitis observed in N20. No acid-fast bacteria were observed in any Ziehl-Neelsen-stained organ or non-cutaneous tissue sections.

**Table 7:** Summary of results and BU status for possums examined during necropsy study.

| Possum ID | Ulcers present? | PCR positive swab(s)? | PCR positive tissue(s)? | Histopathology? | BU status |  |  |
| --- | --- | --- | --- | --- | --- | --- | --- |
|  |  |  |  |  | Clinical BU | Infected | Uninfected |
| N1 | Y – severe | Y | Y | Y – lesions | Y |  |  |
| N2 | N | N<sup>†</sup> | Y | ND |  | Y |  |
| N3 | N | N | Y | ND |  | Y |  |
| N4 | N | N | N<sup>†</sup> | ND |  |  | Y<sup>†</sup> |
| N5 | N | N<sup>†</sup> | Y | ND |  | Y |  |
| N6 | Y – severe | Y | Y | Y – lesions, liver, pancreas | Y |  |  |
| N7 | N | N | N | ND |  |  | Y |
| N8 | Y – moderate | Y | Y | Y – lesions, lungs | Y |  |  |
| N9 | N | N | Y | ND |  | Y |  |
| N10 | Y – mild | N | N | ND |  |  | Y |
| N11 | Y – mild | Y | N<sup>†</sup> | ND | Y |  |  |
| N12 | N | Y | Y | ND |  | Y |  |
| N13 | N | Y | N | ND |  | Y |  |
| N14 | Y – severe | Y | Y | Y – lesions, lungs, liver | Y |  |  |
| N15 | N | Y | Y | ND |  | Y |  |
| N16 | N | Y | N<sup>†</sup> | ND |  | Y |  |
| N17 | N | Y | N<sup>†</sup> | ND |  | Y |  |
| N18 | N | Y | N<sup>†</sup> | ND |  | Y |  |
| N19 | N | Y | Y | ND |  | Y |  |
| N20 (T5) | Y – severe | Y | Y | Y – lesions, liver | Y |  |  |
| TOTALS (Y) – n | 7 | 13<sup>a</sup> | 12<sup>a</sup> | 5 | 6 | 11 | 3<sup>a</sup> |
| TOTALS (Y) – % | 35 | 65<sup>a</sup> | 60<sup>a</sup> | 100 | 30 | 55 | 15<sup>a</sup> |

AFB= acid fast bacteria; BU= Buruli ulcer; ND= not done.
<sup>a</sup> Not including equivocal results.**BU categorisation and geographical distribution of all possums**

#### BU categorisation of the necropsied possums

Of the 20 possums examined during the necropsy study, most (n=11, 55%) were categorised as ‘infected’, 6 (30%) as ‘clinical BU’ and only 3 (15%) as ‘uninfected’, although one of these (N4) had an equivocal PCR result for its spleen sample (see Table 7).

Only one possum that had mild cutaneous lesions suggestive of BU (N10) tested PCR negative for all samples including the swab from its distal tail ulcer, and was therefore ‘uninfected’. A swab from the single small, shallow distal tail ulcer on the other mildly affected possum, N11, also tested PCR negative, but this possum was categorised as ‘infected’ due to its PCR positive cloacal swab (its large intestine result was equivocal).

Of the 11 ‘infected’ possums, oral swabs were positive in 7 (63.6%), and equivocal for an additional 2. The faecal samples from all the ‘infected’ possums were PCR negative, however the cloacal swab was positive for one of these possums (N17), and equivocal for another (N16).

### BU categorisation and geographical distribution of all possums

Combining the data from all 26 possums examined during both the trapping and necropsy studies revealed that 42% (n=11) were ‘infected’, 31% (n=8) were ‘uninfected’ (although one of these possums had an equivocal result), and 27% (n=7) had ‘clinical BU’. The geographical distribution, by BU category, of possums examined in both the trapping and necropsy studies is presented in Fig. 5. The inner Melbourne suburb of Essendon was most strongly associated with ‘clinical BU’, with all four possums examined from this suburb in this category (N1, N14, T5/N20, T6). Most of the possums from the other inner Melbourne suburbs were also ‘clinical BU’ or ‘infected’, with only the possums from Ascot Vale and South Yarra categorized as ‘uninfected’. The Werribee area had a total of three possums examined; two were ‘infected’ and one ‘uninfected’ although this possum (N4) had an equivocal spleen result. Barwon Heads had the lowest association with BU: four possums examined during the trapping study were ‘uninfected’, one trapped possum was ‘clinical BU’ and one necropsied possum that originated from that area was ‘infected’.

**Fig. 5:**
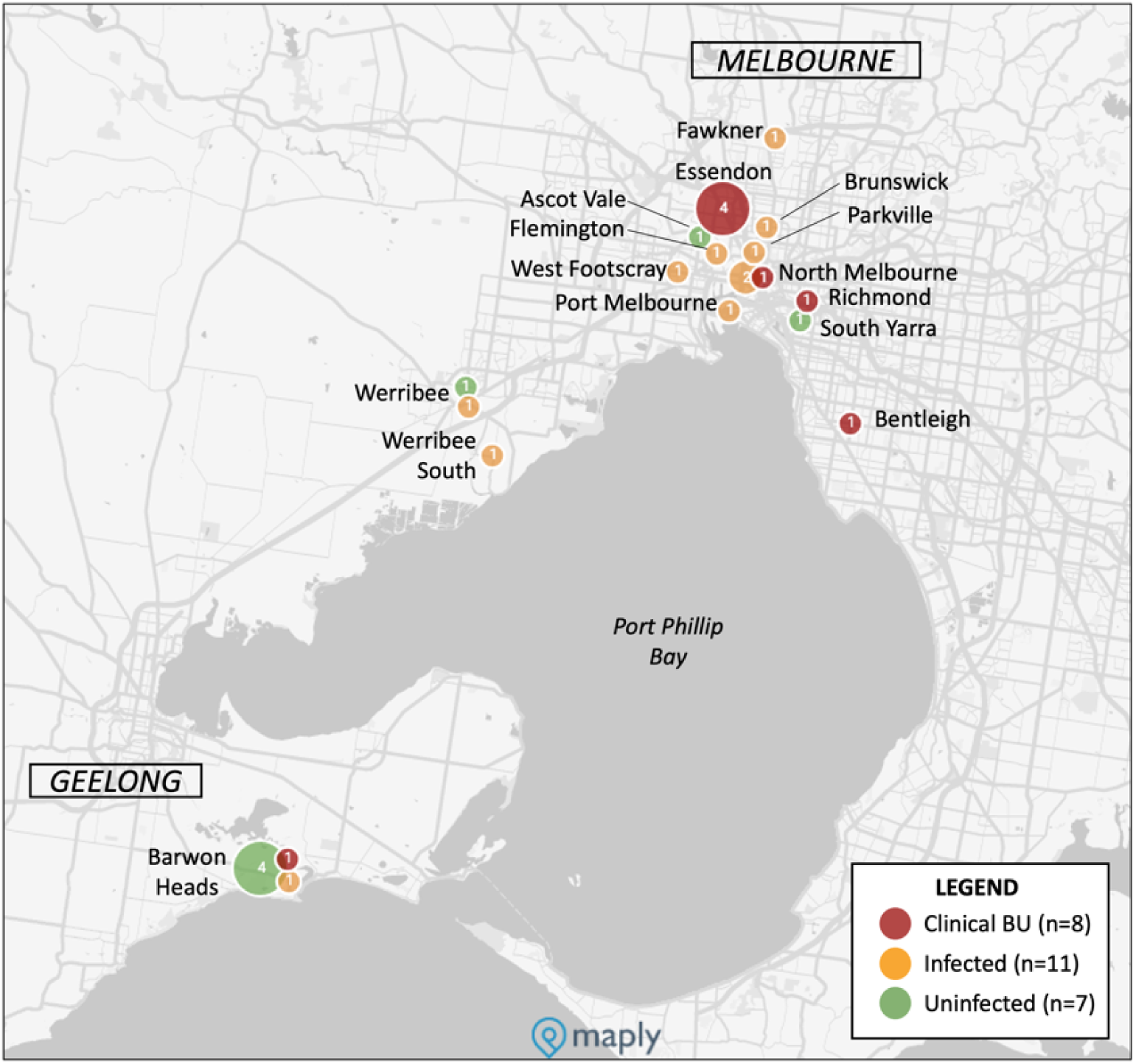
Map of south-central Victoria showing Buruli ulcer (BU) categorization of possums examined in the trapping and necropsy studies. The size of the circle corresponds to the number of possums and the numbers inside the circles state the number of possums.

### Statistical analysis of possum categorical data

Only when comparisons were made between possum species categorised as ‘PCR positive’ (including ‘clinical BU’ and ‘infected’) and those that were PCR negative (‘uninfected’) did a result approach significance (P=0.051), with CRTs having eight times greater odds of being PCR positive than CBTs (see S2.3 Table).

## Discussion

The results presented here confirm the presence of clinical BU and subclinical *M. ulcerans* infection in possums from urban areas of south-eastern Victoria, Australia, including from some previously unreported suburbs in inner Melbourne and Werribee. Our identification of areas of emerging possum BU positivity and the largely negative findings from Barwon Heads, which was a historical hotspot of BU cases in humans and animals in the early- to mid-2000s[16, 19, 22, 42, 43], emphasizes the highly focal and dynamic nature of BU transmission and distribution in Victoria. Surveillance of BU in possums may have public health benefits given that human BU case numbers increase with proximity to *M. ulcerans*-positive possum faeces, and that detection of *M. ulcerans* in possums occurs prior to the emergence of human BU in a geographical region[37, 39]. Our findings provide further evidence that possums are reservoirs of *M. ulcerans* in Victoria[19, 36], and that CRTs are more likely to be PCR positive for *M. ulcerans* than CBTs[36]. The reasons behind this apparent species susceptibility are unclear, but may reflect unique behaviours of CRTs such as caecotrophy (reingesting large moist faecal pellets of high nutritional value originating from the caecum [46, 50], which can selectively retain bacteria in the digestive system for longer periods [51]), and a more social lifestyle that may increase exposure to and transmission of bacteria among individuals, for example while sharing communal tree hollows and dreys[46, 50]. Marsupials, including possums, appear particularly susceptible to mycobacterial infections, potentially due to their unique evolutionary immunobiological pathways[52] and their lower body temperatures compared to placental mammals[53]. Reports of clinical BU in koalas (*Phascolarctos cinereus*) and a long-footed potoroo (*Potorous longipes* from Victoria[19], and detection of *M. ulcerans* DNA in bandicoot (*Isoodon macrourus*) faeces in Queensland[54, 55] support this hypothesis, although systematic testing of Australian wildlife for BU has not been conducted to date.

*M. ulcerans* DNA was detected in 85% of the necropsied possums in this study (17/20, plus an additional ‘uninfected’ possum with an equivocal spleen result), including in every sampled orifice, tissue and organ of the four severely affected possums (N1, N6, N14 and N20/T5) and one moderately affected possum (N8). This confirms that BU is not merely a cutaneous disease in these animals. Systemic distribution of *M. ulcerans* has been previously confirmed in wild CRTs [41, 56] and koalas[20] in Victoria, and in experimental infections of mice and grasscutters (*Thryonomys swinderianus*), with evidence of bacterial circulation in lymph and/or blood[27, 57]. Whether the presence of *M. ulcerans* in the major internal organs is associated with pathology and/or impaired system function in possums has not been established[36, 41], however the moribund condition of two of the severely affected animals in this study indicates that their health was substantially compromised. In contrast, human BU is described almost entirely in skin and subcutaneous tissues, including bones[8, 58], and systemic effects are largely unknown, although secondary bacterial infections have reportedly led to sepsis and other fatal sequelae [59, 60]. There was little consistency in PCR positivity of samples from the ‘infected’ possums, and no clear difference in Ct values in samples from any of the ‘PCR positive’ animals that would suggest amplification of the bacteria occurred in any location. The PCR positivity of certain external tissues including sections of ear, nose, and footpads is interesting, and may be indicative of more widespread dissemination of *M. ulcerans* than was previously expected.

Only those possums with severe and moderate clinical BU lesions had PCR positive faeces, thereby potentially acting as sources of environmental contamination for zoonotic transmission of BU. The faecal samples and cutaneous lesions from the severely and moderately affected necropsy possums and one of the mildly-affected trapped CRTs had the lowest recorded Ct values in our dataset, reflecting highest bacterial loads (Ct values between 14.0 and 20.0 are equivalent to approximately 10^4 to 10^6 *M. ulcerans* genomes[33]). Ulcers and faeces of clinically affected possums are therefore implicated as posing the highest risk of zoonotic transmission to humans, particularly to veterinarians and wildlife handlers who encounter them frequently.

Most of the possums in our study, including those that were ‘infected’, had no skin lesions and had PCR-negative faeces, meaning they would not have posed a transmission risk to humans. This finding, while important and potentially reassuring from a public health perspective, contrasts with other published results in which up to 19% of clinically unaffected possums did have PCR positive faeces[19, 36]. O’Brien, Handasyde (36) also demonstrated that the faecal positivity of sub-clinically affected possums can change over time, although the underlying drivers for this changing status have not been elucidated. Our finding that one of the trapped CBTs (T4) had negative oral, cloacal and ulcer swabs but positive faeces is interesting and difficult to explain. Longitudinal trapping studies are needed to further investigate the factors influencing faecal shedding of *M. ulcerans* in possums.

Our necropsy study findings that 71% (12/17) of ‘PCR positive’ possums had positive oral swabs (plus 2 with equivocal swab results) strongly supports the hypothesis that possums are ingesting *M. ulcerans,* presumably from a yet-unidentified[19, 34] environmental source. CRTs are almost exclusively arboreal[46] and are therefore unlikely to routinely encounter *M. ulcerans* that may be present in soil or water bodies. Targeted PCR sampling of the native trees, flowers and fruit that CRTs are known to prefer may provide crucial missing data on the environmental source(s) of infection in this species. Possums could also ingest *M. ulcerans* while licking their cutaneous ulcers, however this would not explain the 64% oral swab positivity observed in the ‘infected’, clinically unaffected possums (7/11, plus 2 equivocal) unless they were regularly engaging in communal grooming, which is not a recognised behaviour in this species ([46, 61], personal observations, ECH, PW).

Other potential mechanisms of *M. ulcerans* infection or transmission in possums may be suggested by our results. Mapping of the cutaneous lesions from the seven clinically affected possums revealed that distal limbs (paws) and tails were the most frequently affected regions, which supports earlier findings[19, 36]. Paws and tails are used for climbing and may more frequently incur puncturing injuries and lacerations that facilitate invasion of *M. ulcerans* from the environment, although this is perhaps less likely to explain lesions on the dorsal surfaces of paws. Distal limbs, facial regions and ventral tail surfaces are relatively less furred than other parts of the body and may be more accessible biting sites for mosquitoes. The two *M. ulcerans*-positive mosquitoes collected during the Essendon trapping event supports this theory. Blood meal analyses have shown that *Ae. notoscriptus* readily feed on humans and native possums in Victoria [33, 62], and mapping of BU lesions in humans revealed a similar pattern in that distal limbs – areas of exposed skin typically not covered by clothing in summer and which are preferred biting sites for certain mosquitoes – were the most frequently affected body regions[63]. Possums’ facial regions may more frequently incur injuries that are susceptible to bacterial invasion during fighting; a behaviour that is typically more common in males[46, 61] and which may provide an explanation for the previously reported increased risk of clinical BU in male CRTs [36]. The observed positivity of pouch swabs in our study may suggest that juvenile possums – highly altricial and weighing less than 1 gram at birth[61] – are exposed to *M. ulcerans* throughout their entire early stages of development. Our finding that all six juvenile (back joey) possums examined via necropsy were either ‘infected’ (n=5) or ‘clinical BU’ (n=1) may lend some support to this theory. The proposed transmission cycle of *M. ulcerans* in Victoria is depicted in Fig. 6.

**Fig. 6:**
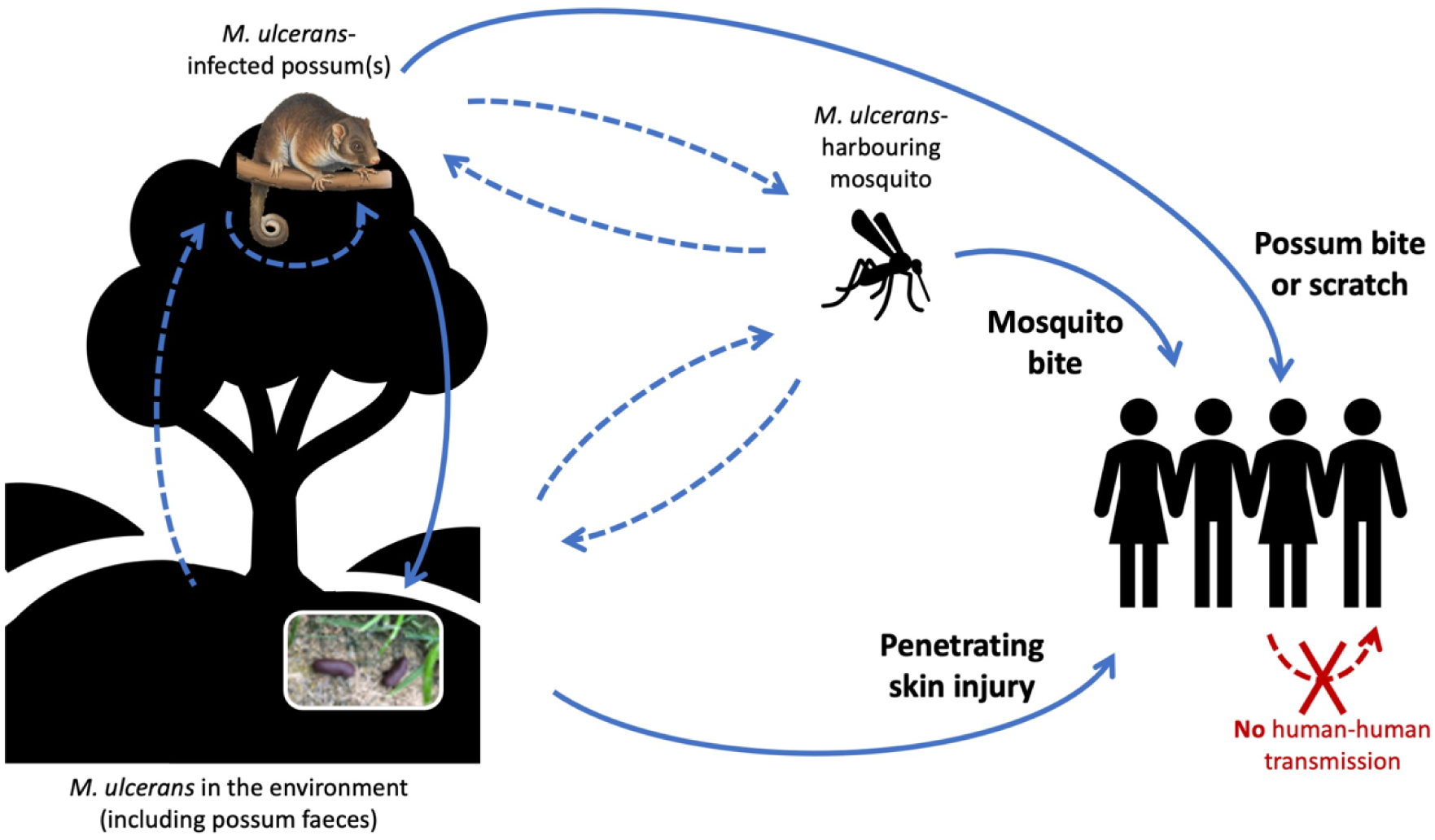
Schematic of proposed *Mycobacterium ulcerans* transmission cycle in Victoria, Australia, showing mosquitoes as mechanical vectors, possums as wildlife reservoirs, and humans as incidental hosts. Solid blue lines represent known transmission pathways; dashed blue lines represent presumed but unknown pathways. Possums may be acquiring *M. ulcerans* via numerous means: from the environment, via ingestion of an unidentified source or by contamination of wounds incurred during climbing or fighting; from bites of *M. ulcerans*- harbouring mosquitoes; or from other possums, either vertically, being exposed to *M. ulcerans* during early development in the pouch, or horizontally, via exposure to *M. ulcerans* shed in possum faeces.

Our study has limitations. Possum cadavers were sourced from two veterinary clinics, which would likely have biased collection of possum cases from certain suburbs. Both clinics frequently receive wildlife case referrals from neighbouring areas, however, which may have mitigated that bias to some extent. Informing the clinical staff that possums were being collected for a BU study may have made them more likely to remember to retain cadavers that had cutaneous lesions. Repeated longitudinal trapping events that were planned at Barwon Heads as part of this study were not able to be conducted within the project timeframe; consequently, trapped possums were only sampled at one point in time. While our findings provide important additions to our understanding of BU in Victorian possums, more research is required to estimate BU prevalence and incidence, and to establish the geographical distribution of BU in possums in Victoria. Limitations also exist with regards to the interpretation of PCR results. Several samples returned Ct values above 40, which was the defined threshold for PCR positivity, and some other samples tested in duplicate returned equivocal results, preventing clear interpretation of infection status. Although these samples were not classified as positive, the high Ct values may reflect the presence of low levels of *M. ulcerans* DNA associated with early or late infection. Similarly, possum BU categorisation was determined by PCR positivity, and one female CRT (N4) was categorised as ‘uninfected’ despite an equivocal PCR result from her spleen sample. Finally, despite following standard necropsy protocols it is possible that some cross-contamination of *M. ulcerans* may have occurred during tissue dissection, which could have artificially inflated the number of PCR positive samples from individual possums.

The evolving ecology and unique transmission dynamics of BU in Victoria provide ongoing challenges for disease control, and an integrated One Health approach involving further research, targeted interventions and educational outreach is needed. BU is not a notifiable animal disease in any Australian state or territory, and awareness of BU is typically low among veterinarians and wildlife clinics (personal observations, ECH, PW), even in highly endemic areas such as the Mornington Peninsula, south-east of Melbourne in Victoria, where human BU case incidence is among the highest in the world[16, 64]. Targeted dissemination of educational information would increase awareness of BU in native wildlife and the potential risk of zoonotic transmission to humans, especially those in high-risk locations and occupations. Animal BU testing has only been conducted as part of specific research projects to date. Structured veterinary BU surveillance is encouraged and may identify novel areas of disease foci, which would also have important implications for public health given the reported spillover of BU to humans subsequent to detection in possums[39]. Sampling natural predators of CRTs, including dogs, foxes, goshawks, powerful owls and wedge-tailed eagles[65], may also shed light on how

*M. ulcerans* is introduced into new geographical areas. A recent study detected *M. ulcerans* DNA and RNA in a small number of fox faecal samples, suggesting that foxes can excrete live and presumably infectious bacteria[34]. Given the distribution of foxes throughout urban Australia, and their large home ranges – up to 150 hectares in parts of Victoria[66] – it is plausible that foxes may ingest infected CRTs in one location and subsequently excrete viable *M. ulcerans* in new, previously BU-negative areas. This warrants further research. While PCR testing enables accurate diagnosis, it may not be readily available in all regions. Development of *M. ulcerans*-specific point-of-care diagnostic tests such as loop-mediated isothermal amplification (LAMP) and recombinase polymerase amplification (RPA) assays[67–69] are promising, but would need robust validation and assessment studies to determine test accuracy under field conditions and on animal-derived samples.

Finally, the involvement of mosquitoes and possums in transmission of *M. ulcerans* in Victoria provides opportunities for One Health-based disease control interventions. Reducing populations of *Ae. notoscriptus* is possible via combinations of chemical- and community-based mosquito control methods, however sustained delivery over the entire Victorian endemic area would be labour- and resource-intensive, and success may be hampered by acceptability and uptake of interventions by affected communities[70]. Possum-based interventions would have the dual benefits of improving animal health and welfare, and protecting public health. Intensive antibiotic treatment regimes, frequent dressing changes and surgical skin grafting are typically curative in humans[47] but unlikely to be feasible for wild animals. At present, severely afflicted possums presented to veterinary clinics are typically euthanised, but widespread culling of possum populations for disease control purposes would be neither legal, ethical nor effective: many studies evaluating the effects of wildlife culling on infectious disease dynamics have found counterproductive increases in disease transmission and geographical expansion of pathogen distribution[71–73]. Oral bait vaccination of possums against BU is a potential control option. Oral bacille Calmette-Guérin (BCG) vaccines are currently used in New Zealand for control of *Myocobactierum bovis* in introduced Australian CBTs, to protect the public from zoonotic tuberculosis[74, 75], and early laboratory studies have shown that BCG vaccines provide cross-protection against *M. ulcerans* infections in mice[76]. Research is needed to determine the optimal formulation and delivery methods for distribution and uptake of effective and palatable vaccines, tailored to the different possum species residing in BU-endemic areas[77].

Human BU case numbers and geographical distribution are increasing in Victoria and this trend is expected to continue, especially given the effects of climate change already being experienced across much of the Victorian BU-endemic area: mild winters, heavy rainfall and floods are associated with higher CRT survival rates [65] and increased mosquito breeding[78, 79]. A One Health approach is needed to control transmission of *M. ulcerans* in Victoria, to protect both animal and public health from this important emerging disease.

## Conclusion

Cases of *M. ulcerans* infections and clinical BU in native possums in Victoria – particularly CRTs – may be more numerous and widespread than previously believed. BU can cause severe cutaneous disease that substantially impairs the health and welfare of affected possums, and systemic internal distribution of the bacteria may be associated with additional health impacts. Infected possums may pose a risk of zoonotic transmission to humans, particularly to those in high-risk occupations such as veterinarians and wildlife handlers, noting that infection requires inoculation of the bacteria into subcutaneous tissue. Possums may be contracting BU via numerous pathways, including via bites of *M. ulcerans*-harbouring mosquitoes. Given their involvement in zoonotic transmission of *M. ulcerans* in Victoria, possum-based disease surveillance and management interventions could have benefits to public health. Potential opportunities include ongoing possum excreta surveys to identify areas with increased transmission risk, enabling more targeted mosquito control and improving community education. An integrated One Health approach is needed for successful BU control in this region.

## Acknowledgements

The authors wish to acknowledge the work of the ‘Beating Buruli in Victoria’ team members and contributors, including Peter Mee, Andrew Buultjens, Ary Hoffman, Véronique Paris, Nick Bell, Stacey Lynch, Nick Golding, David Price, Koen Vandellanoote, Jane Oliver and Paul Johnson.

We would like to thank Tristan Rich, Samantha Lovett and the staff of Lort Smith Animal Hospital, the veterinary staff of the former U-Vet Werribee Animal Hospital, Jessica Haining and other Anatomic Pathology lab staff at the Melbourne Veterinary School who provided invaluable assistance with our necropsy study, and Faye Docherty and Mirjana Bogeski for histology and histochemistry. We are also grateful to Ian Beveridge, Mark Stevenson, Liz Dobson, Richard Ploeg, Smitha Georgy and Christina McCowan for generously providing their parasitological, statistical and pathological expertise.

We would also like to thank Jasmin Hufschmid, Kath Handasyde, Carolyn O’Brien, Alisdair Legione, Brett Gardner, Sarah Frith, Peter Smolderen, Katherine Whittaker and Anna-Katarina Schilling for their valuable advice and assistance with the possum trapping study. Big thanks also to Maddie Glynn, Jon Paskas and the staff at the Barwon Heads Caravan Park for all their assistance and support, and to Natalie J. Russell for her tireless advocacy on behalf of the Essendon possums.

## Supporting information

**S1 Appendix. Possum examination sheets.**

S1.1 Examination record sheet for adult and sub-adult possums examined during the trapping studies.

S1.2 Examination record sheet for back young examined during the trapping studies.

S1.3 Final examination record sheet for possums necropsied during the study. The study protocol and record sheet were slightly amended to add some tissues (heart, brain and bone marrow/femur) after the study had commenced.

**S2 Appendix. Additional results tables.**

S2.1 Table. IS2404 PCR results from mosquitoes opportunistically collected during Essendon possum trapping, December 2022.

S2.2 Table. IS2404 PCR result outcomes for swabs and tissue samples collected during the possum necropsy study.

S2.3 Table. Statistical comparison of possums with at least one PCR positive sample (‘clinical BU’ and ‘infected’) and with no PCR positive samples (‘uninfected’).

